# p75 Neurotrophin Receptor Shapes the Dynamics of Adult Hippocampal Neurogenesis in Alzheimer’s Disease

**DOI:** 10.1101/2025.09.11.675537

**Authors:** Maria Anna Papadopoulou, Konstantina Chanoumidou, Maria Peteinareli, Electra Tsaglioti, Konstantina Michalaki, Matthieu D. Lavigne, Ioannis Charalampopoulos

## Abstract

Adult hippocampal neurogenesis is essential for cognitive flexibility and emotional resilience, and its disruption is strongly associated with Alzheimer’s disease, a disorder marked by cognitive decline and memory impairment. The p75 neurotrophin receptor regulates neuronal survival and plasticity, yet its contribution to adult hippocampal neurogenesis, especially under neurodegeneration, remains unclear. In this study, we investigate the role of p75NTR in Adult Neurogenesis using constitutive and conditional p75NTR knockout mice, the amyloidogenic 5xFAD model, and 5xFAD/p75NTR knockout mutants. We show that p75NTR deletion led to a significant reduction in NSC proliferation and altered neuronal differentiation in the dentate gyrus, acting in a cell non-autonomous function to control neural stem cell fate. Notably, 5xFAD/p75NTR mutants displayed exacerbated neurogenic deficits compared to 5xFAD mice. Transcriptomic profiling confirmed these alterations and supported a disease-relevant regulatory function. Parallel studies in human iPSC-derived neural stem cells exposed to amyloid-β showed p75NTR-dependent mechanisms mirroring findings from the mouse models. Collectively, our findings establish p75NTR as a critical regulator of adult hippocampal neurogenesis, under Alzheimer’s Disease and propose it as a therapeutic target.

## INTRODUCTION

Adult neurogenesis is the process of continuous generation of new neurons by the proliferation and selective differentiation of endogenous Neural Stem Cells (NSCs) throughout life, specifically in the adult hippocampus, a region critical for cognitive functions like learning and memory. This age-dependent, dynamic process involves the proliferation, migration, differentiation and proper integration of newborn neurons into existing neural circuits, contributing to cognitive flexibility and emotional resilience (Dumitru et al., 2025; Kempermann, 2016; Ming & Song, 2011; C. Zhao, 2006). Noticeably, adult neurogenesis is a scientific field with many contradictory findings, and mounting evidence implicates impaired adult hippocampal neurogenesis in the cognitive decline observed in Alzheimer’s Disease (AD), highlighting the significance of elucidating the molecular mechanisms underlying this phenomenon (Flor-García et al., 2020; Moreno-Jiménez et al., 2019; Salta et al., 2023).

AD, the most common form of all dementias, consists one of the most pressing challenges in healthcare system, characterized by progressive cognitive decline, memory impairment, and ultimately, profound disability (A. Armstrong, 2019). Despite decades of research, effective disease-modifying treatments for AD remain elusive, underscoring the urgent need for novel therapeutic approaches. Among the various pathological hallmarks of AD, including extracellular amyloid-beta (Aβ) plaques, intracellular neurofibrillary tangles (NFTs) and synaptic dysfunction, recent emerging evidence suggests that dysregulation of adult hippocampal neurogenesis plays a pivotal role in disease onset and progression (Armstrong, 2009; Mu & Gage, 2011). Under AD, the hippocampal region is severely damaged and is linked to reduced adult neurogenesis and memory deficits, in both humans and animal models (Jahn, 2013; Moreno-Jiménez et al., 2019; Mu & Gage, 2011).

Neurotrophins and their receptors, key molecules for the development and functional sustainment of the nervous system, are expressed in various stem cell types and play a pivotal regulatory role with respect to stem cell differentiation, survival and migration (Chao, 2003; Pramanik et al., 2017). They are secreted growth factors, namely NGF, BDNF and NT3/4, whose main functions are to induce neuronal survival and regeneration (Huang & Reichardt, 2001), by activating the high-affinity TrkA, TrkB or TrkC receptors respectively, while all mature and immature neurotrophins activate p75 pan-neurotrophin receptor (p75NTR) (Barnabé-Heider & Miller, 2003; Dechant & Barde, 2002). Noticeably, neurotrophic factors control the adult postnatal neurogenesis in the main neurogenic regions of the brain, meaning the Sub-Granular Zone (SGZ) of the hippocampus and the Sub-Ventricular Zone (SVZ) of the lateral ventricles (Shohayeb et al., 2018; Vilar & Mira, 2016).

The pan-neurotrophin p75 receptor, a member of the Tumor Necrosis Factors Receptors (TNFRs) superfamily, is being widely expressed in most cell types among the neural tissue, throughout the developing brain. However, it becomes refined to specific brain regions during adulthood and is essential for neuronal cell death and/or cell survival, differentiation and synaptic plasticity (Chao, 2003; DeFreitas et al., 2001; Lewin & Carter, 2014; Nykjaer & Willnow, 2012). The p75NTR exhibits diverse functions depending on its cellular context, interacting partners, and the availability of its ligands, in both developmental and adult stages (Charalampopoulos et al., 2012; Ibáñez & Simi, 2012; Vilar et al., 2009). As a multifactorial receptor, p75NTR, is gaining interest as a “fate decision protein” in stem cells, modulating their potency and differentiation (Tomellini et al., 2014). Several studies have been initiated with the aim to describe p75NTR-dependent actions on NSCs, since it is emerging as a key regulator of adult neurogenesis, although with contradictory results that rather confuse the field than define the role of the receptor (Bernabeu & Longo, 2010; Boskovic et al., 2014; Catts et al., 2008; Dokter et al., 2015; Poser et al., 2015), due to the diverse genetic backgrounds of p75NTR-deficient animals and species-specific variations.

Given the critical role of adult hippocampal neurogenesis in maintaining cognitive function, receptor’s disruption has increasingly been recognized as a contributing factor to neurodegenerative disorders, particularly AD (Zeng et al., 2011). p75NTR is highly expressed in degenerative cell populations in AD and has been also extensively linked to Amyloid-β (Aβ) -the major component of the plaques-pathological signaling, by serving as a receptor for this molecule, either by promoting neuronal cell death or by protecting against Aβ-induced toxicity (Bengoechea et al., 2009; Saadipour et al., 2013; Yao et al., 2015). Thus, it is evident that the precise mechanisms underlying p75 receptor-mediated regulation of adult hippocampal neurogenesis in the context of AD, remain poorly understood.

Based on our previous expertise with small-molecule neurotrophin mimetics [reviewed in (Zota et al., 2025)], we have shown that BNN27, a synthetic derivative of the neurosteroid DHEA, selectively targets TrkA and p75NTR receptors and exerts multimodal neuroprotective effects. In the 5xFAD mouse model of AD, chronic BNN27 administration promoted proliferation, counteracted Aβ-induced cytotoxicity, and regulated astroglial inflammatory responses through p75NTR-dependent pathways. This work underscored the therapeutic potential of p75NTR modulation in AD, while also highlighting key molecular targets and mechanisms underlying adult neurogenesis deficits in AD (Kokkali et al., 2024).

In the present study, we sought to elucidate the role of the p75NTR in adult hippocampal neurogenesis using *in vivo* mouse models and human NSCs. We delineate the specific p75NTR signaling pathways which regulate NSC proliferation and differentiation under AD, by using genetically modified mouse models of three different ages (2, 4 and 6 months old): i) the 5xFAD model which is a well-established AD animal model that resembles many of the amyloidogenic pathology of AD (Oakley et al., 2006), ii) the p75NTR fully knock out mouse p75NTRExIII model (firstly described by (Lee et al., 1992)), iii)the cross breeding between these two models, creating the 5xFAD/p75NTR ko mouse model, as well as iv) the p75 fl/fl NestinCre mouse, in order to study the cell-autonomous effects of the receptor upon its specific deletion in NSCs. Furthermore, we have recently shown the receptor’s expression in human induced Pluripotent Stem Cell (hiPSC)-derived NSCs (Papadopoulou et al., 2023). Thus, we now investigate p75NTR role in proliferation of human NSCs (hNSCs) as a human model to recapitulate Amyloid beta (Αβ)-associated toxicity. Finally, we are using an RNA sequencing based high-throughput strategy to evaluate the genetic differential expression analysis of samples derived by 2 months old p75NTR ko, 5xFAD and 5xFAD/p75NTR ko mice. Our observations clearly indicate that the presence of p75NTR is an essential factor for regulating neural stem cell properties, and our results provide for the first time a detailed tempospatial description of the p75NTR-mediated effects in adult neurogenesis under physiological and neurodegenerative conditions, implying p75NTR as a connective molecule for the neurogenic deficits on AD, and thus as a novel target of pharmacological manipulation.

## RESULTS

### p75NTR is a key regulator of NSC proliferation

To test the effect of p75NTR deficiency on the proliferation of NSCs in adult mice, we compared the number of BrdU and Sox_2_ positive cells at the DG of the hippocampus in WT and p75NTR ko mice. The deletion of p75NTR in the ko mice model was identified by the absent expression of the receptor in the Basal Forebrain after immunohistochemical staining using primary antibody for p75NTR (Fig. EV1).

Coronal sections of hippocampal DG from WT and p75NTR ko mice were co-immunostained with BrdU and Sox_2_ primary antibodies and analysis showed that the number of proliferative NSCs was significantly decreased in the p75NTR ko mice [from 1011.25 ± 66.3 cells (SEM) to 530.5 ± 63.6 cells (SEM)] (Fig. 1A, B), corresponding to a 47.5% decrease. Thus, these results indicate that the expression of p75NTR is necessary for the proper proliferation of NSCs in the adult mice of 2 months old. If we take into consideration 4- and 6-months old mice, we observed that the deficiency of p75NTR in 4-months old mice, still has a significant effect on the proliferation rates reducing their numbers from 765 ± 38.2 cells (SEM) in WT to 489.5 ± 35 cells (SEM) in p75NTR ko mice (Fig. 1A, B). On the other hand, in 6 months old mice there were no significant differences since the proliferation of NSCs diminishes to such an extent (approximately around 75%) at this age that no significant effect in proliferation can be detected.

**Fig. 1.**
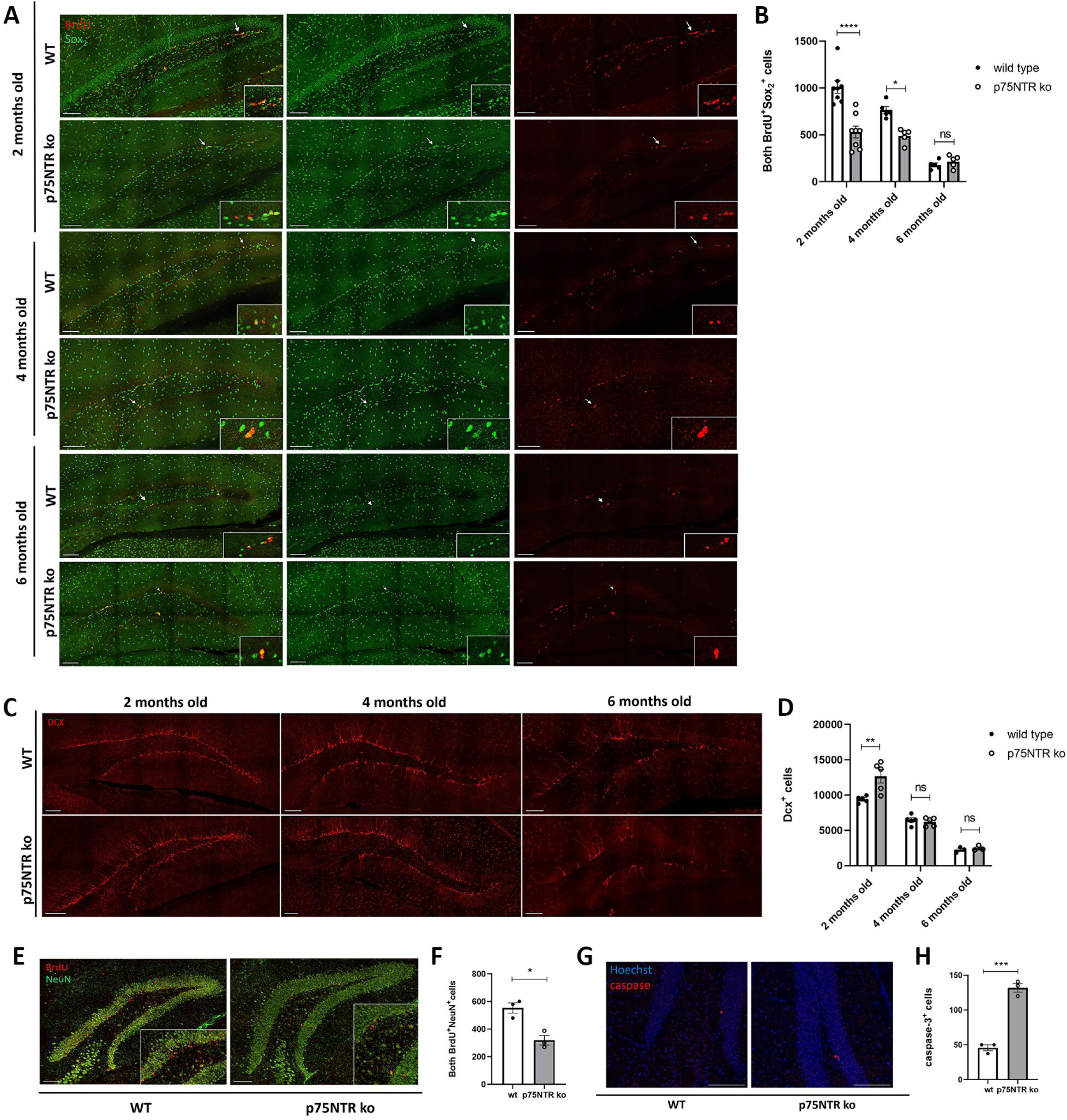
The effects of p75NTR deficiency on the proliferation, differentiation and maturation of NSCs. (A) Coronal sections, of the hippocampal DG from 2 months old WT and p75NTR ko mice injected with BrdU for 5 days. Sections were co-immunostained for BrdU (red) and Sox_2_ (green). Scale bar, 100 μm. (B) Quantification of both BrdU^+^ and Sox_2_^+^ cells in injected mice (2mo -- n=8 for each genotype; 4mo & 6mo -- n=5 for each genotype). Data are presented as mean ± SEM. 2way ANOVA ****p<0,0001, *p<0,0,5 ns, no significant. (C) Coronal sections, of the hippocampal DG from 2 months old WT & p75NTR ko mice. Images depict Dcx (red) immunostained immature neurons. Scale bar, 100 μm. (D) Quantification of Dcx^+^ cells in p75NTR ko & WT mice (2mo, 4mo & 6mo -- n=5 for each genotype). Data are presented as mean ± SEM. 2way ANOVA**p<0,005, ns, no significant. (E) Coronal sections, of the hippocampal DG from 2 months old WT mouse. Images depict BrdU (red) & NeuN (green) immunostained mature neurons. Scale bar, 100 μm. (F) Quantification of both positive cells in 2 months old p75NTR ko & WT mice (n=3 p75NTR ko, n=3 WT). Data are presented as mean ± SEM. *p<0.05 (Student’s t-test unpaired). (G) Coronal sections of the hippocampal DG from 2 months old WT mouse. Images depict caspase3^+^ cells (red), Hoechst (blue). Scale bar, 100 μm. (H) Quantification of caspase3^+^cells in 2 months old WT & p75NTR ko mice (n=3 p75NTR ko, n=3 WT). Data are presented as mean ± SEM. ***p<0.001 (Student’s t-test unpaired). BrdU, 5-bromo-2′-deoxyuridine; Sox_2_, SRY-Box Transcription Factor 2; Dcx, doublecortin; NeuN, Neuronal Nuclear marker.

### Impairment of neuronal maturation in p75NTR deficient mice

To test the effect of p75NTR deficiency on the differentiation of NSCs of the hippocampal DG towards the production of immature neurons at the age group of 2, 4 and 6 months old, we studied the number of Doublecortin (Dcx) positive cells, a protein marker depicting immature neurons. To our surprise, the analysis showed significantly increased number of immature neurons in the p75NTR ko mice compared to the WT [from 9410 ± 207 cells (SEM) to 1267.6 ± 949.7 cells (SEM) in p75NTR ko mice] (Fig. 1C, D). Furthermore, the area of Dcx^+^ cells’ processes was also significantly increased in p75NTR ko mice (Fig. EV2). Thus, the reduction of proliferation that was observed in 2 months old p75NTR ko mice resulted in the significant increase of immature neurons, indicating a robust maturation process forced by the lack of p75NTR expression. This receptor is known to regulate cell cycle progress (Underwood & Coulson, 2008; Vilar et al., 2006) and this action could explain the increased maturation rates in NSCs. Moreover, we studied the effect of p75NTR deletion in NSC differentiation in 4- and 6-months old mice (Fig. 1C, D), where we found no significant differences. As expected, the number of Dcx^+^ cells in 6 months old mice was reduced compared to this of 2 months old mice (approximately around 75%) following the same pattern with the proliferation procedure.

### Reduction of neuronal maturation in the p75NTR ko mice

To further evaluate the hypothesis of p75NTR-mediated cell cycle deterioration, driving immature neurons to “freezing”, we also investigated the production of fully mature neurons at 2 months old mice. For that reason, we pulsed mice with BrdU for the first 5 days of a 3-week period and after 21 days overall we sacrificed the mice and studied the number of BrdU^+^ and NeuN^+^ (Neuronal nuclear marker of mature neurons) cells that survived. Our analysis showed that the number of both positive cells in the DG was significantly decreased in p75NTR ko mice compared to the WT [from 553.3 ± 37.2 cells (SEM) to 319.17 ± 35.1 cells (SEM) in p75NTR ko mice] (Fig. 1E, F). Thus, p75NTR is necessary not only for proper differentiation of neuronal precursors to immature neurons, but it also determines the final number of fully mature neurons. In order to specify if this decreased number of NeuN^+^ cells is due to the inability of Dcx^+^ neurons to further differentiate or cells are more vulnerable, we measured cell death in hippocampal DG. We stained and counted the caspase-3 positive cells, and we revealed that there is a significantly increased number of cell death under p75NTR deletion [from 45.83 ± 4.1 cells (SEM) to 131.87 ± 6.15 cells (SEM) in p75NTR ko mice] (Fig. 1G, H). Contradictory to receptor’s well known pro-apoptotic effects, we here show that its expression is important for the proper differentiation and maturation of NSCs, by sustaining their survival rates in addition to cell cycle control.

### Genes differential expression analysis confirmed the neurogenic capacity of p75NTR

To identify the gene networks underlying the role of p75NTR in adult neurogenesis, we performed RNA sequencing on hippocampal tissue obtained from WT mice, p75NTR ko, and a newly generated mouse line upon crossing of 5xFAD with the p75NTR ko mice, the 5xFAD/p75NTR ko (with the WT group serving as the reference). This bioinformatic analysis revealed 9 significantly upregulated and 33 significantly downregulated genes, as visualized in the volcano plot (Fig. 2A). Gene Ontology (GO) and pathway enrichment analyses revealed that these differentially expressed genes (DEGs) are significantly involved in biological processes such as the regulation of neurogenesis, axonogenesis, cell growth, and cell migration (Fig. 2B), confirming the biological results from the *in vivo* studies. Genes and GOs are detailed in Data Set EV1. A heatmap representing the expression patterns of all 42 DEGs is shown in Fig. 2C, highlighting several genes with known roles in proliferation and migration. For instance, *Med1* is involved in the induction of cell proliferation and migration and regulates BDNF levels (Nagpal et al., 2018), while *Msl1* contributes to cell cycle progression (X. Li et al., 2009). *Fabp7* promotes proliferation and survival via the MEK/ERK signaling pathway (Ma et al., 2018), and deletion of *Fxr1* has been shown to impair neurogenesis by reducing cell proliferation in the dentate gyrus (Patzlaff et al., 2017). Additionally, *Cdh1* plays a critical role in regulating neurogenesis and cortical development (Delgado-Esteban et al., 2013), and, together with *Fzr*, promote neural stem cell differentiation in Drosophila (Ly & Wang, 2020). Conversely, genes such as *Sv2a* were identified for their role in supporting NSC survival through the p53 signaling pathway (Yu et al., 2023).

**Fig. 2.**
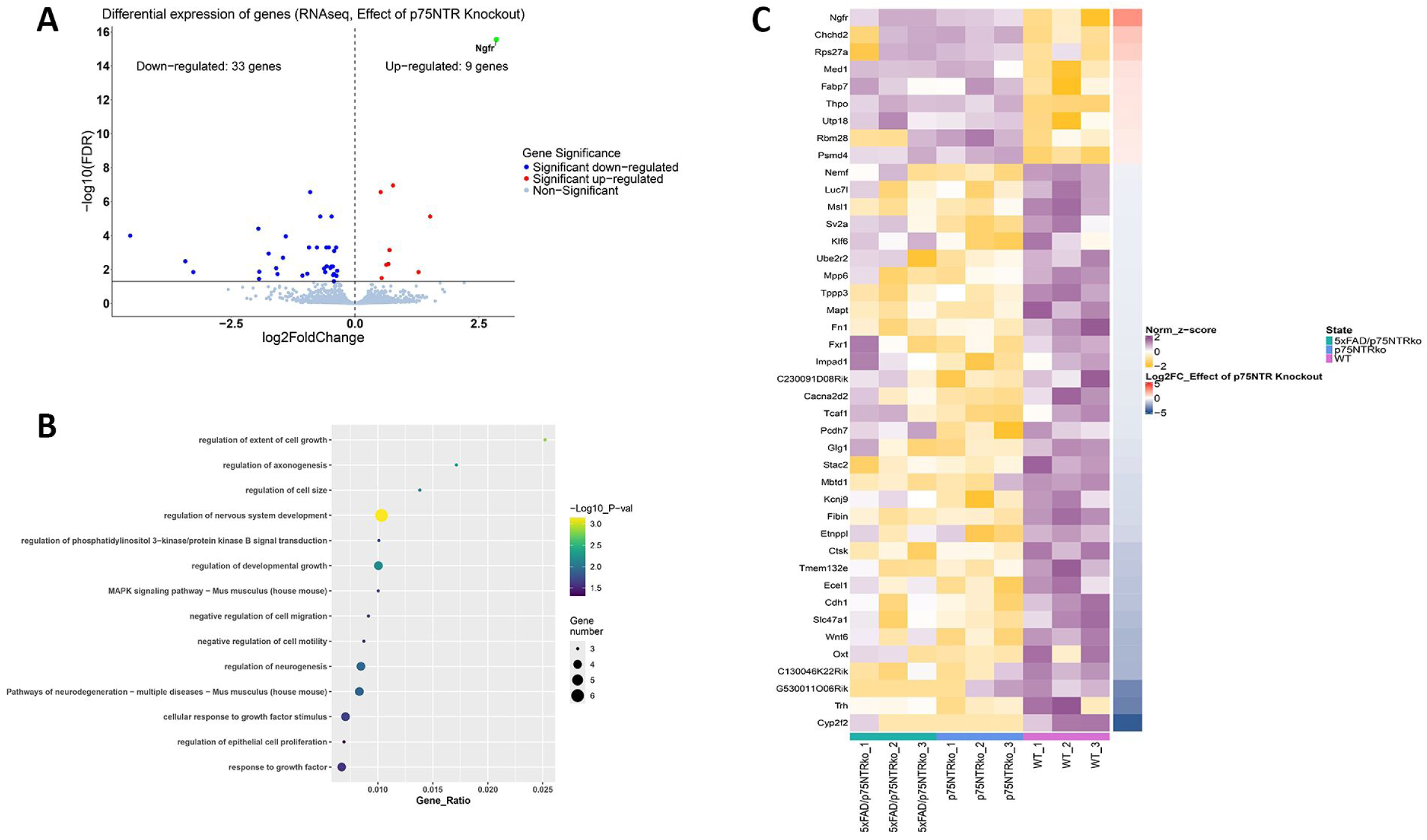
Functional analysis of differential gene expression between p75NTR ko mice and 5xFAD/p75NTR ko, with healthy WT mice treated as the reference. (A) Volcano plot showing the number of Up (9) & Down (33) regulated genes (p adj < 0.05), pointing out the differential expression of *Ngfr*. (B) Bubble plot showing selected gene ontology (GO) and pathways significantly enriched with differentially expressed genes (DEGs) discovered. Analysis was performed using Metascape (https://metascape.org). The x axis represents the gene ratio. Size of the bubble represents the numbers of DEGs found in each GO list. Coloration from yellow to black represents the – Log10_P value and depicts low (yellow) to high (black) enrichment scores (p adj < 0.05). (C) Heatmap showing the expression profile of all differentially expressed genes (p adj < 0.05) between samples with (5xFAD/p75NTR ko - green, p75NTR ko - blue) and without (WT - purple) the p75NTR receptor knocked out. Scaling was performed across genes using z-scores, to highlight relative expression differences. Fold changes are represented using the log₂ fold change (log₂FC) scale to reflect differential expression between conditions.

It is of special notice that the *Ngfr* gene was found to be upregulated in both p75NTR ko and 5xFAD/p75NTR ko mice (Fig. 2A, C), indicating the important functions of this receptor since its expression levels are induced as a result of its loss of function. It could also be suggested that the receptor’s intracellular part remaining after exon III deletion has potential signaling properties that contribute or opposing its extracellular functions. This upregulation was validated via qRT-PCR (Fig. EV3), with results presented as logarithmic fold change (logFC) relative to WT controls. To further explore the biological implications of these transcriptomic changes, we performed Gene Set Enrichment Analysis (GSEA), which revealed enrichment of GO Biological Process categories such as neural precursor cell proliferation, regulation of neuronal differentiation, and regulation of apoptotic signaling in p75NTR ko mice compared to WT (Fig. EV4, Data Set EV1).

### p75NTR is affecting NSC proliferation in a cell non autonomous mechanism

In order to validate whether p75NTR expression specifically in NSCs is majorly controlling the adult neurogenesis in the DG or if there are other cellular mechanisms derived from p75NTR expression in other cell types, like astrocytes, oligodendrocytes or microglia, we used an animal model, namely p75 fl/fl Nestin Cre, by crossing mice with a specific deletion of p75NTR exonII with the Nestin-Cre mice, in order to delete the receptor only in Nestin positive cells (since Nestin is expressed only in NSCs). We counted again both BrdU^+^Sox_2_^+^ cells, at the DG of the hippocampus and the analysis showed that the number of proliferating NSCs in the DG remained unchanged in p75 fl/fl NestinCre mice compared to the WT (Fig. 3A, C). Thus, p75NTR is affecting adult neurogenesis in a cell non autonomous mechanism.

**Fig. 3.**
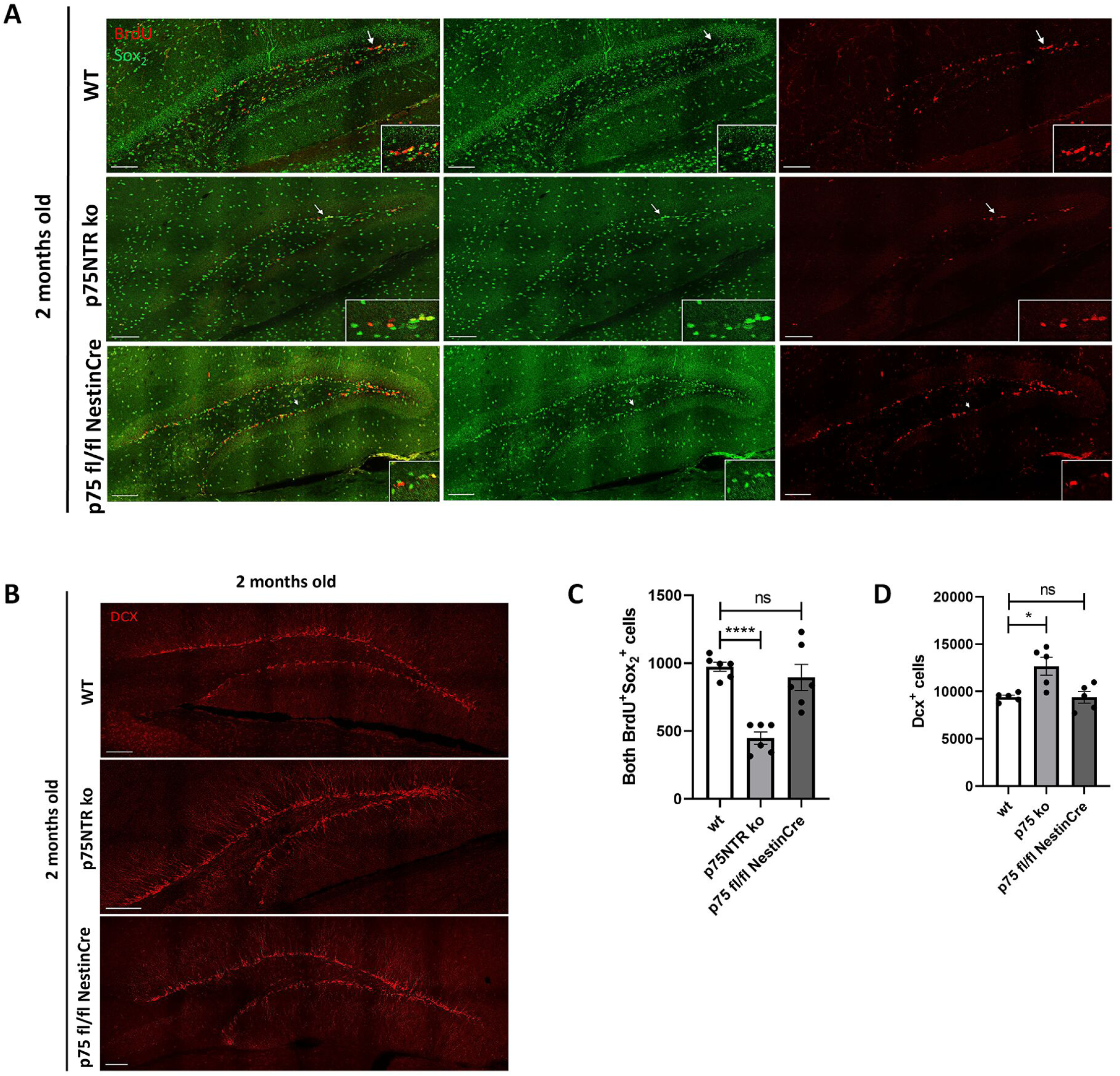
p75NTR deficiency in p75 floxed/floxed NestinCre mice is not affecting the proliferation and differentiation of NSCs. (A) Coronal sections of the hippocampal DG from 2 months old WT and p75 fl/fl NestinCre mice injected with BrdU for 5 days. Sections were co-immunostained for BrdU (red) and Sox_2_ (green). Scale bar, 100 μm (B) Coronal sections of the hippocampal DG from 2 months old WT & p75 fl/fl NestinCre mice. Images depict Dcx (red) immunostained immature neurons. Scale bar, 100 μm. (C) Quantification of BrdU^+^ and Sox2^+^ cells in injected mice (n = 6 for each genotype). Data are presented as mean ± SEM. p**** < 0,0001, ns, no significant (one way ANOVA). (D) Quantification of Dcx^+^ cells in 2 months old WT & and p75 fl/fl NestinCre mice (n=5 for each genotype). Data are presented as mean ± SEM. *p < 0,05, ns, no significant (one way ANOVA). BrdU, 5-bromo-2′-deoxyuridine; Sox_2_, SRY-Box Transcription Factor 2; Dcx, doublecortin.

### p75NTR expressed in Nestin^+^ cells does not affect the production of immature neurons

Furthermore, we analyzed if p75NTR specifically expressed in Nestin^+^ cells has an impact in the production of immature neurons. Thus, we counted the number of Dcx^+^ cells in the hippocampal DG of 2 months old p75 fl/fl NestinCre mice. Our results showed that there were no significant differences between the production of immature neurons in WT and p75 fl/fl NestinCre mice (Fig. 3B, D). In agreement with the aforementioned data about the proliferation of NSCs in this mouse model, we suggest that p75NTR expressed specifically in NSCs does not influence their proliferation, neither the production of immature neurons, clearly indicating a cell non autonomous mechanism of action in adult neurogenesis.

### Increased proliferation rates of NSCs in 2 months old 5xFAD mice

In order to study the proliferation of NSCs under neurodegenerative conditions such as AD, we performed immunofluorescent staining of the coronal sections of hippocampal DG of 2 months old 5xFAD mice. Initially, we tested 5xFAD homozygous *vs* heterozygous mice to choose which genotype is better suited to our experiments. There were no significant differences between 2-months old 5xFAD heterozygous and homozygous mice (Appendix Figure S1). Thus, we decided to continue our studies with the 5xFAD heterozygous mice, since they are also more commonly used in research studies. As it is shown in Fig. 4A and 4B, there is a statistically significant increase in the proliferation rates of NSCs in 5xFAD mice when compared to the WT [from 997 ± 28.2 cells (SEM) to 1271.15 ± 80.8 cells (SEM) in 5xFAD mice]. Next, we studied the number of proliferative NSCs in 4 months old 5xFAD mice, and we observed that there is a significant decrease in the number of both BrdU^+^ and Sox_2_^+^ cells [from 765 ± 38.2 cells (SEM) to 475.5 ± 38.4 cells (SEM) in 5xFAD mice], as expected, since at this age most of the neurodegenerative effects of AD-like pathology have been described to be on their onset in this mouse strain (Fig. 4A, B). Additionally, 6 months old 5xFAD mice showed no significant differences, since in that age the number of proliferative NSCs is very low (Fig. 4A, B). The aforementioned results clearly show that AD background induces hippocampal neurogenesis, at the initial stages at least. We thus hypothesize that the increased number of NSCs observed in the 5xFAD background may represent a compensatory, homeostatic response to the neurodegenerative effects of AD-like pathology, such as the specific result of increased expression of Aβ, which could enhance signaling neurogenic mechanisms (J. Kim et al., 2007; Lopez-Toledano, 2004; Wang et al., 2016; Whitson et al., 1990). This response could potentially aim to counteract the neuronal loss typically seen at the onset of the disease, which generally occurs around 4 months of age (Kokkali et al., 2024).

**Fig. 4.**
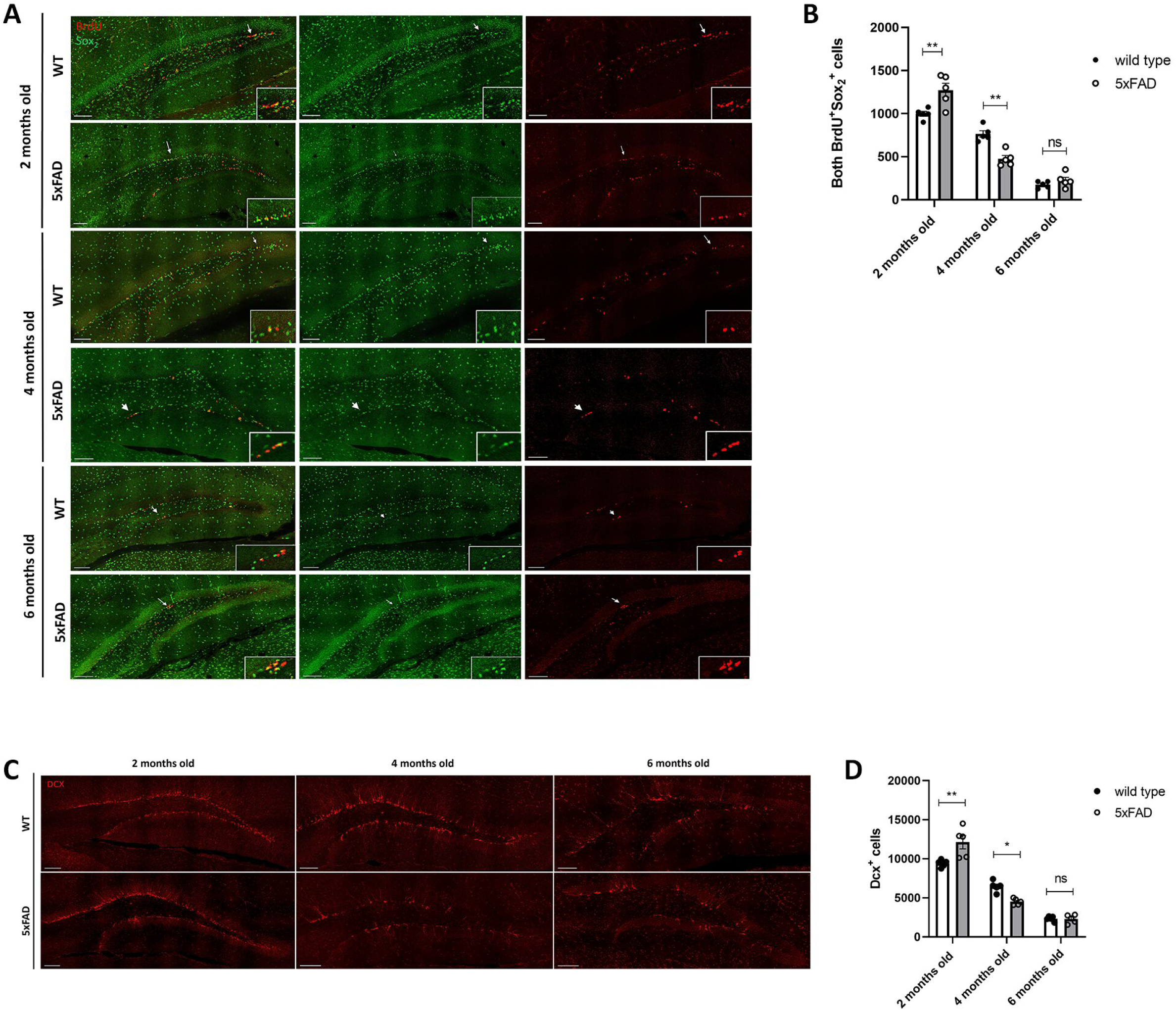
Proliferation and differentiation of NSCs, under Alzheimer’s Disease. (A) Coronal sections, of the hippocampal DG from 2 months old WT and 5xFAD mice injected with BrdU for 5 days. Sections were co-immunostained for BrdU (red) and Sox_2_ (green). Scale bar, 100 μm. (B) Quantification of BrdU^+^ and Sox_2_^+^ cells in injected mice (2mo, 4mo & 6mo -- n=5 for each genotype). Data are presented as mean ± SEM. 2way ANOVA **p<0,005, ns, no significant. (C) Coronal sections, of the hippocampal DG from 2 months old WT & 5xFAD mice. Images depict Dcx (red) immunostained immature neurons. Scale bar, 100 μm. (D) Quantification of Dcx^+^ cells in WT & 5xFAD mice (2mo, 4mo & 6mo -- n=5 for each genotype). Data are presented as mean ± SEM. 2way ANOVA**p<0,005, *p<0,05, ns, no significant. BrdU, 5-bromo-2′-deoxyuridine; Sox_2_, SRY-Box Transcription Factor 2; Dcx, doublecortin.

### 2 months old 5xFAD mice present increased number of immature neurons

In order to assess the production of immature neurons in terms of AD, we measured the Dcx^+^ cells in 2 months old 5xFAD mice. By comparing the Dcx^+^ cells in the DG of the hippocampus of WT and 5xFAD mice, we concluded that there is an increase, with a significant difference between the two groups [from 9410 ± 210 cells (SEM) to 12128.25 ± 847.8 cells (SEM) in 5xFAD mice] (Fig. 4C, D). These results confirm that AD phenotype affects the proliferation of NSCs and, in continuation, has also a significant impact in the production of new immature neurons, further proposing a general regulatory mechanism of neurogenesis upon Aβ changes. Taking into consideration, the number of immature neurons in 4 months old 5xFAD mice, we observed a significant decrease in the number of Dcx^+^ cells [from 6469.6 ± 310.2 cells (SEM) to 4511.2 ± 180.6 cells (SEM) in 5xFAD mice], as it was expected due to the progress of the neurodegenerative effects of AD (Fig. 4C, D). As it was previously shown in WT mice, the 6 months old 5xFAD mice had no differences, emphasizing on the really low number of immature neurons at this age (Fig. 4C, D).

### Deficiency of p75NTR in 2 months old 5xFAD mice reversed the increased proliferation of NSCs

Based on the above results, we tested the outcome of p75NTR deletion in the 5xFAD mice. We generated double transgenic mice, the 5xFAD/p75NTR ko and by following the same experimental protocols and measuring the number of both BrdU^+^Sox_2_^+^ cells, we revealed that in 2 months old 5xFAD/p75NTR ko mice there was a significant decrease in the proliferation rates of NSCs when compared to the WT mice, almost at the same levels as in p75NTR ko mice [from 997 ± 28.2 cells (SEM) to 702 ± 40.5 cells (SEM) in 5xFAD/p75NTR ko mice] (Fig. 5A, B). This novel result confirms the crucial role of p75NTR for the proliferation of NSCs, even in the AD background. It is of special notice also that this p75NTR-mediated inhibition of NSC proliferation counteracts the 5xFAD genotype, where an elevated NSC proliferation was observed. Considering the results derived by the 4 months old mice, we observed for once more, a decreased number of proliferative NSCs in the 5xFAD/p75NTR ko mice, showing again the importance of p75NTR expression [from 765 ± 38.2 cells (SEM) to 176 ± 17.7 cells (SEM) in 5xFAD/p75NTR ko mice] (Fig. 5A, B). Furthermore, if we compare the number of proliferative NSCs in the 5xFAD and 5xFAD/p75NTR ko mice, of both ages 2- and 4-months old, we can conclude that the expression of p75NTR is necessary for the proliferation of NSCs under AD both during the onset as well as the progression of the disease (Fig. 5A, B).

**Fig. 5.**
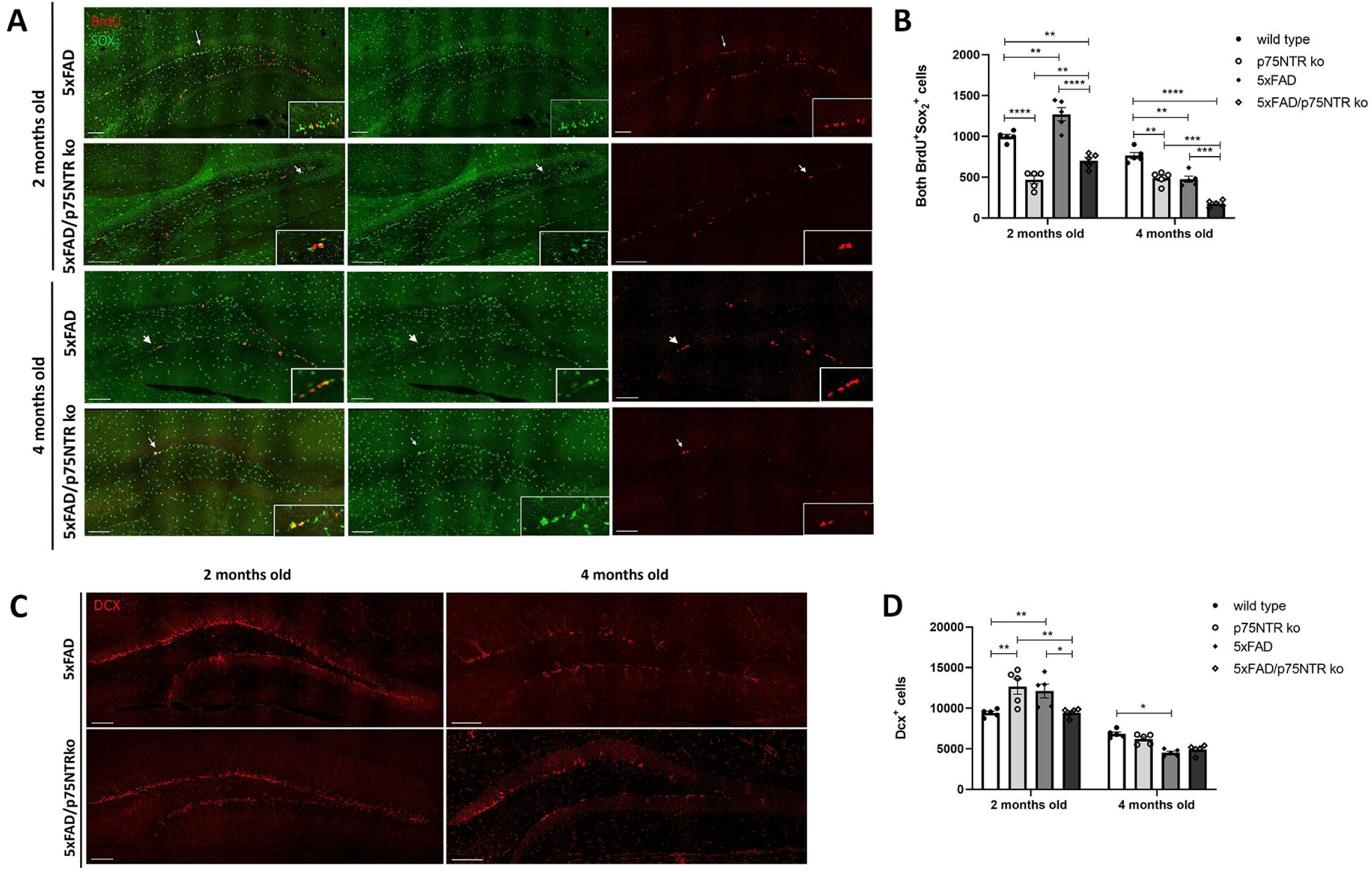
The effects of p75NTR deficiency on the proliferation and differentiation of NSCs, under Alzheimer’s Disease. (A) Coronal sections, of the hippocampal DG from 2 months old 5xFAD/p75NTR ko mouse injected with BrdU for 5 days. Sections were co-immunostained for BrdU (red) and Sox_2_ (green). Scale bar, 100 μm (B) Quantification of BrdU^+^ and Sox_2_^+^ cells in injected mice. (2mo & 4mo - n=5 for each genotype). Data are presented as mean ± SEM. 2way ANOVA****p<0,0001, ***p<0,001, p**< 0,005. (C) Coronal sections, of the hippocampal DG from 2 months old WT, p75NTR ko, 5xFAD and 5xFAD/p75NTR ko mice. Image depicts Dcx (red) immunostained immature neurons. Scale bar, 100 μm. (D) Quantification of Dcx^+^ cells in WT, p75NTR ko, 5xFAD, and 5xFAD/p75NTR ko mice (2mo & 4mo - n=5 for each genotype). Data are presented as mean ± SEM. 2way ANOVA**p<0,005, *p<0,05. BrdU, 5-bromo-2′-deoxyuridine; Sox_2_, SRY-Box Transcription Factor 2; Dcx, doublecortin.

### p75NTR deficiency in 2 months old 5xFAD mice does not affect the production of immature neurons

Having the above in mind, we analyzed the number of Dcx^+^ cells upon p75NTR deletion in 2 months old 5xFAD mice. The number of Dcx^+^ cells in the DG of the hippocampus of 5xFAD/p75NTR ko mice was reduced compared to p75NTR ko and the 5xFAD mice, and was almost the same as this of the WT mice of the same age (Fig. 5C, D). So, p75NTR deletion is sufficient to abolish the increase in NSC proliferation rates of 5xFAD and also to affect the production of immature neurons and decrease the number of Dcx^+^ cells, compared to p75NTR ko and 5xFAD mice [from 12128.25 ± 847.8 cells (SEM) in 5xFAD mice to 9448.25 ± 243.2 cells (SEM) in 5xFAD/p75NTR ko mice] (Fig. 5C, D).

Comparing the number of Dcx^+^ cells, in 4 months old mice, we observed that although there is a decreased number of immature neurons in the 5xFAD/p75NTR ko mice, this number is not statistically different from that one of the 5xFAD model (Fig. 5C, D). Taking into consideration the significantly reduced number of Dcx^+^ cells at the age of 4 months old, we assume that any potential effects of receptor’s deletion are minimized and cannot be detected, if there are any.

### Gene Set Enrichment Analysis revealed genes implicated in p75NTR-dependent effects under AD

To quantitatively evaluate the gene networks associated with the role of p75NTR in AD, we performed RNA sequencing on hippocampal tissue obtained from 5xFAD and 5xFAD/p75NTR ko mice. This analysis identified five significantly upregulated and one downregulated gene, as illustrated in the volcano plot (Fig. EV5).

Among the upregulated genes, *Wdfy1*, which interacts with TLR4, is known to promote neurogenesis and facilitate the recruitment of TLR3 and TLR4 signaling adaptors that mediate neural stem cell differentiation and the production of type I interferons and inflammatory cytokines (Hu et al., 2015). Thbs4, another upregulated gene, is implicated in the migration of newly formed neurons from the rostral migratory stream (RMS) to the olfactory bulb (Girard et al., 2014). In contrast, the downregulated gene *Hes5* plays a critical role in maintaining NSCs, regulating neuronal differentiation, and controlling the timing of the transition between neurogenesis and gliogenesis during mammalian neocortical development (Bansod et al., 2017). To further investigate the functional relevance of p75NTR in the context of AD, we conducted GSEA which revealed that pathways related to neurogenesis, cellular differentiation, and Wnt signaling were enriched in 5xFAD/p75NTR ko samples, whereas pathways associated with cell death, inflammation, DNA damage, stress responses, and regulation of lipid biosynthesis were predominantly enriched in 5xFAD samples (Fig. EV5). Genes are detailed in Data Set EV2.

### p75NTR expression in hiPSCs-derived NSCs

Although the use of humanized mouse models as tools for studying human diseases is still meaningful, the translational outcome of any research work demands testing in human tissue or cells. To fulfil this approach, we have explored the p75NTR-dependent effects on neurogenesis using human iPSCs-derived NSCs. This methodology allows also the validation of mouse results, and provides a novel platform for drug screening of new compounds with neuroprotective and/or neurogenic effects against AD. We have generated NSCs from two different iPSC lines, derived from healthy individuals (named as 841, 856). To study the role of p75NTR in human NSCs, we firstly detected its expression in NSC lysates with Western Blot analysis. Apart from p75NTR, NSCs were found to express the p75NTR intracellular interactors RIP2 and TRAF6. Furthermore, the actual interaction of p75NTR with the aforementioned TRAF6 protein also confirmed with co-IP (Fig. 6A) indicating an active state of p75NTR signaling in these cells.

**Fig. 6.**
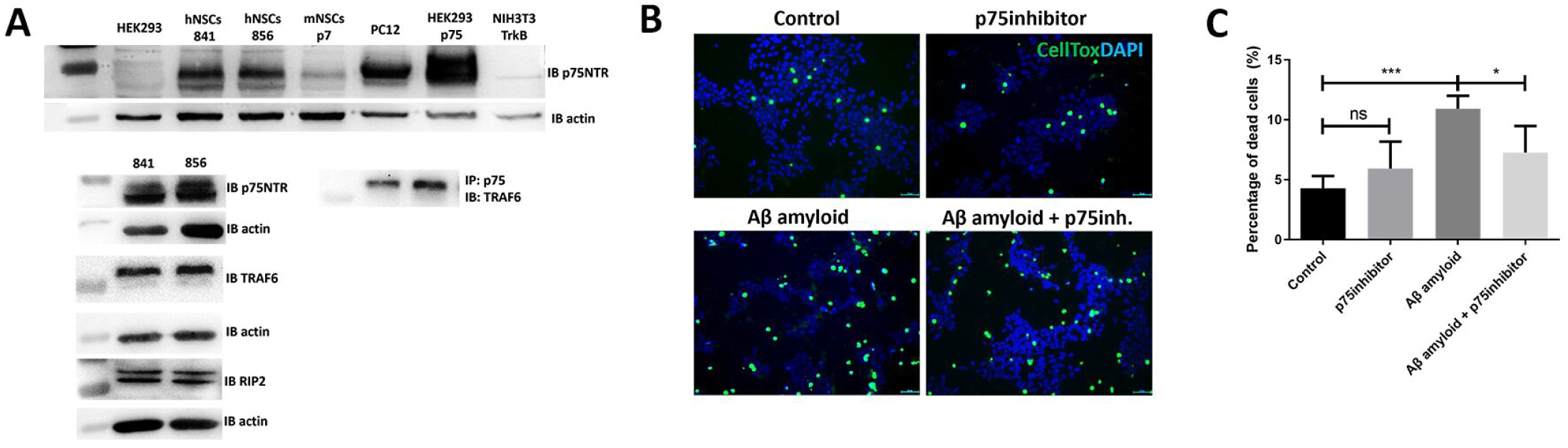
p75NTR expression and function on human iPSCs - derived NSCs. (A) Representative blots (n=3) determined via Western Blot showing the expression of p75NTR, RIP2 and TRAF6 proteins in lysates of two iPSCs-derived NSCs lines (841, 856). Co-IP showing the interaction of p75NTR with TRAF6 protein. HEK293T cells, mNSCs p7 (mouse NSCs from postnatal day 7), PC12 (pheochromocytoma of the rat adrenal medulla), HEK293T cells transiently transfected with p75NTR plasmid and NIH3T3 cells expressing TrkB receptor were used like control samples. (B) Dead cells (green) upon inhibition of p75NTR and/or treatment with aβ oligomers (50μm scale bar). (C) Graph showing the percentage of dead cells (Celltox labeled cells/total number of HOECHST^+^ cells) n=3 from different NSCs lines (841, 856) (unpaired t-test ***p < 0,0001, *p < 0,05, ns, no significant).

### Inhibition of p75NTR rescues Aβ induced toxicity in human iPSCs – derived NSCs

Following the detection and signaling properties of p75NTR, we were interested in investigating the necessity of p75NTR on NSC survival under healthy and AD relevant conditions. To that end, we blocked the activity of p75NTR by using a p75NTR specific neutralizing antibody (2,5ngr/ml, MC-192) which has been extensively used as a selective pharmacological inhibitor of p75NTR activation. AD pathology was mimicked by treating the cells with Aβ peptides (specifically the Aβ1-42 oligomers) that are known to have a toxic effect on cells. The results of cell death measurement using the Celltox assay, provide clear evidence that p75NTR negatively influences human NSC survival after treatment with Aβ peptides indicating a regulatory role of p75NTR in human NSC pathology of AD (Fig. 6B, C).

### Inhibition of p75NTR has no effect in proliferation of human NSCs

In continuation of the p75NTR-mediated cell survival effects, we also investigated the effect of p75NTR inhibition on NSC proliferation under healthy conditions. BrdU assay in NSCs, revealed no alteration in the proliferation rate of the cells when p75NTR is inhibited, suggesting that p75NTR is not actively involved at least in human NSC proliferation (Appendix Figure S2). This finding is in agreement with our results in mice studies, since it proves the cell non autonomous effects of this receptor to NSC proliferation.

## DISCUSSION

Alzheimer’s Disease, the most challenging neurodegenerative disease in terms of epidemiological penetrance and socioeconomical cost, is characterized by the extensive, non-reversable, neuronal loss, which finally leads to incapability to sustain life. There is no effective treatment for AD at the moment and numerous clinical trials of potential medications that target tau or amyloid dysfunctions have been merely successful (Anand et al., 2014; Kodamullil et al., 2017). Recent novel studies have targeted more or even new aspects of the disease, including neuroinflammation, autophagy and neurogenesis in order to offer multimodal therapeutic advances (Chen et al., 2024; Tang et al., 2024; R. Zhao, 2024). In addition to targeting the later stages of disease progression, earlier events—such as the reduction of neuroprotective molecules like neurotrophins—may also represent valuable pharmacological targets for promoting brain repair.

Neurotrophins and their receptors consist an important class of endogenous molecules for development and maintenance of nervous system, especially under pathological challenges. Their pleiotropic signaling through Trk and p75NTR receptors orchestrates multicellular signals to promote neuronal repair and regeneration (Chao, 2003; Huang & Reichardt, 2001; Pramanik et al., 2017). Enabling selective activation of neurotrophin receptors provides strong pharmacological tools on the armamentarium against neurodegenerative diseases (Kokkali et al., 2024; Nagahara & Tuszynski, 2011; Tuszynski, 2024; Yi et al., 2021), since slowing or blocking disease progression and ultimately increasing neurogenesis to restore neuronal loss would be an effective therapeutic strategy (Magavi et al., 2000; Mu & Gage, 2011).

In the present study, we explored the interconnected triangle, p75NTR-AD-adult hippocampal neurogenesis, aiming to decipher their interplay. p75 neurotrophin receptor has been recently emerged as a key molecule for mediating anti-Alzheimeric protection (Shanks et al., 2024) and adult neurogenesis enhancement, while its high affinity ligand, proNGF acts as specific mitogen factor in NSCs (Corvaglia et al., 2019). To this scope, we used p75NTR ko mice to study the receptor’s effects under physiological conditions.

As depicted in Figure 1, deletion of p75NTR resulted in reduced NSC proliferation, increased number of immature Dcx^+^ cells and decreased number of mature NeuN^+^ cells, indicating p75NTR-dependent cell cycle arrest. Indeed, it has been shown that p75NTR regulates the cell cycle and facilitates the cell cycle exit of neuronal cells (Underwood & Coulson, 2008; Vilar et al., 2006). Interestingly, p75NTR loss of function induced cell death in the hippocampal region, controversial to receptor’s major function as a pro-apoptotic mediator.

p75NTR has been shown to be expressed in the DG and studies revealed its expression by progenitor cells located in the SGZ and SVZ (Barrett et al., 2016; Bernabeu & Longo, 2010; Catts et al., 2008; Colditz et al., 2010; Young et al., 2007). To further delineate p75NTR activity in NSC properties, we generated a mouse line by crossing p75NTR floxed mice with the Nestin-Cre mice, in order to delete receptor’s expression only in Nestin^+^ cells, meaning the NSC population in the adult brain. Measuring proliferation and early differentiation in these mice, we did not observe any significant changes in the number of proliferating NSCs and also in the number of immature neurons compared to the WT mice (Fig. 3). Thus, it seems that p75NTR being expressed by NSCs, has no role in adult neurogenesis and differentiation of these cells, and we assume that there is a non-autonomous mechanism that is being activated. According to literature, p75NTR is not only expressed by NSCs but also by other cell types like astrocytes and oligodendrocytes (Becker et al., 2018; Cragnolini & Friedman, 2008). Even if its expression is restricted to a small population in the adult rodent and human CNS (Dowling et al., 1999), after different types of injury or disease, astrocytes proliferate and there is a strong induction of p75NTR expression (Oderfeld-Nowak et al., 2003; Qin et al., 2022) as well as by oligodendrocytes (Tep et al., 2013; Zota et al., 2024). Thus, p75NTR combinatorial activity in some of these cell types could contribute in a cell non-autonomous manner to neural stem cell proliferation and differentiation.

Another interesting finding of the present study was the upregulation of the remaining intracellular part of 75NTR. By performing RNA-seq analysis, we revealed signaling pathways and genes that are related to neurogenesis processes, but most importantly we observed in our surprise that the most prominent effect was in *Ngfr,* which is actually the deleted gene. To confirm this finding, we examined *Ngfr* levels by quantitative RT-PCR. It is known that the p75NTR ko mouse line that was used in the present study (deletion of exonIII) can still express the short p75NTR isoform (von Schack et al., 2001). The observed increase of this remaining, uncapable of neurotrophin-dependent signaling, intracellular domain could be explained by a feedback loop of the cellular system to regain p75NTR functions, indicating its importance in homeostasis. Although, the truncated receptor is not able of mediating neurotrophin actions, the existing protein could mimic naturally occurring proteolytic fragments of p75NTR and being capable of pro-apoptotic signaling.

Upon characterization of the role of p75NTR in adult neurogenesis under physiological conditions, we investigated its impact under neurodegenerative pathology, such as in AD. For this purpose, we generated 5xFAD mice that lack the expression of p75NTR, the 5xFAD/p75NTR ko mice. As controls, we used not only the WT and p75NTR ko mice, but also the 5xFAD mice, where we measured in 2- and 4-months old mice the proliferation and differentiation in the DG. We show that 2 months old 5xFAD mice have increased number of proliferative and differentiative NSCs (Fig. 4), an effect that could be explained as a compensatory mechanism or as a selective signaling property of the overproduced Amyloid-beta (Aβ) which characterizes this strain. Even though Aβ peptides are considered neurotoxic, they can mediate many biological processes, both in adult brains and throughout brain development. Lopez-Toledano showed that neurogenesis is induced by Aβ42 and it increases the total number of neurons *in vitro* in a dose-dependent manner (Lopez-Toledano, 2004) while Aβ peptide in its monomeric form and at low concentrations may be neuroprotective, and enhances the survival of hippocampal neurons *in vitro* (J. Kim et al., 2007; Wang et al., 2016; Whitson et al., 1990). In agreement with this hypothesis, at 4 months old 5xFAD mice both proliferating and differentiating cells are decreased compared to WT, a result that follows the well-established knowledge of the toxic effects of Aβ in NSCs. We should highlight here that oligomerization and deposition of Aβ in the 5xFAD mice begins after 2 months (Oakley et al., 2006), while at 4 months old, 5xFAD mice start to develop plaques, whereas plenty of Aβ deposits could be detected in 8-months old 5xFAD mice (Ziegler-Waldkirch et al., 2018).

Having described the NSCs stages at the 5xFAD mice, we finally investigated the proliferation and differentiation of NSCs in the 5xFAD/p75NTR ko mice. Deficiency of p75NTR in 2 months old 5xFAD mice reversed the increased proliferation of NSCs that we observed in 5xFAD mouse model (Fig. 5A, B). Thus, it seems that p75NTR’s deletion is overcoming genotype in order to abolish the high proliferation of the 5xFAD model. The levels of Dcx^+^ cells follow the same pattern, and if we compare these mice with their control groups, meaning 5xFAD model and p75NTR ko, we observe a significant reduction of immature neurons (Fig. 5C, D). Thus, it seems that the p75NTR deficiency in an AD background reversed the AD-dependent high rates of proliferative NSCs. If we take into consideration all of the above information, we could propose that in the 5xFAD mice, where p75NTR expression is present, Aβ could probably interact with the receptor, leading to neuroprotection (Bengoechea et al., 2009; Saadipour et al., 2013; Yao et al., 2015). However, in 5xFAD/p75NTR ko mice, Aβ cannot act through p75NTR and no neuroprotective effects are induced, resulting on the decreased number of proliferative NSCs and Dcx^+^ cells.

5xFAD/p75NTR ko mice of 4mo, showed a really decreased number of proliferative NSCs, showing that the deficiency of p75NTR and the degenerative effects of AD which arise in 4mo mice, are both strong enough to drop the levels of the number of proliferative NSCs at these mice (Fig. 5A, B). Furthermore, if we compare the number of proliferative NSCs in the 5xFAD/p75NTR ko model and in the 5xFAD mice, we observe again a really decreased number of BrdU and Sox_2_ positive cells, showing this accumulating negative effect (Fig. 5A, B).

Comparing the number of Dcx^+^ cells, in 4 months old 5xFAD/p75NTR ko mice, we observed it is not statistically different from that one of the 5xFAD mice nor the WT (Fig. 5C, D). It seems that in 4mo the deficiency of p75NTR has no impact in the differentiation of these cells, as it does earlier at the age of 2mo. All the results from our *in vivo* experiments are summarized in Table 3.

In order to validate the results from our animal models to a more translational outcome, we generated human NSCs produced by hiPSCs, expressing p75NTR (Fig. 6A). Then, we evaluated the downstream mediators of the p75NTR signaling pathways, RIP2 and TRAF6 intracellular proteins, in human NSCs, showing an active state of p75NTR signaling in these cells. Furthermore, we show here, that p75NTR negatively influences human NSC survival after treatment with Aβ amyloid (Fig. 6B, C) indicating a regulatory role of p75NTR in NSC pathology of AD. Our findings reveal the involvement of p75NTR signaling in NSCs, and our upcoming research will focus on a comprehensive characterization and detailed investigation of its role in human neurogenesis in AD patients.

Overall, the findings of the present study pave the way to decipher the specific signaling properties of p75NTR in adult neurogenesis under physiological and neurodegenerative conditions, as AD. By deciphering the exact properties and functions of the p75NTR in NSC proliferation, differentiation and integration, as well as on the other cell populations, we could pharmacologically target this receptor in order to develop novel therapeutic strategies against neurodegeneration. By validating the results from *in vivo* mouse studies in humanized models which resemble the cell properties as well as the connectivity of neuronal networks, we could provide more robust translational platforms for drug screening and thus accelerate therapeutic interventions for neurological disorders.

## MATERIALS AND METHODS

### Mice

Wild-type (WT), p75NTR ko, p75 fl/fl NestinCre, 5xFAD and 5xFAD/p75NTR ko male and female mice were used of different ages (two, four and six-months old). p75NTR ko mice (Ngfrtm1Jae targeted mutation 1, Rudolf Jaenisch p75NTR/ExonIII-) were obtained from the Jackson Laboratory, Bar Harbor, ME (Strain #:002213) and maintained on C57BL/6 background. The p75fl/fl mice with the loxP sites targeting the ExonII of p75NTR gene were kindly provided by Dr Sebastian Thieme from the Technical University of Dresden (Ngfrtm1a(EUCOMM)Wtsi) and crossed to transgenic mice containing the Nestin gene driving expression of Cre recombinase. 5xFAD heterozygous mice were obtained from the Jackson Laboratory (#034848-JAX) and maintained on C57BL/6 background. 5xFAD mice express human APP and PSEN1 transgenes with a total of five AD-linked mutations, the Swedish (K670N/M671L), Florida (I716V), and London (V717I) mutations in APP, and the M146L and L286V mutations in PSEN1 (Oakley et al., 2006). 5xFAD/p75NTR ko were generated by crossing 5xFAD heterozygous with p75NTR heterozygous mice. All mouse models were kept in the Animal House of the Institute of Molecular Biology and Biotechnology (IMBB-FORTH, Heraklion, Greece), in a temperature-controlled facility on a 12 h light/dark cycle, fed by standard chow diet and water ad libitum. Animal experimentation received the approval of Veterinary Directorate of Prefecture of Heraklion, Crete and was carried out in compliance with Greek Government guidelines and the guidelines of FORTH ethics committee.

### BrdU labelling

BrdU i.p. injections 100 mg/kg (10mgr/ml-Sigma, St. Louis, MO, USA, B5002) were performed for 5 days (1 injection per day). Analysis was performed either immediately after the BrdU injections (assessment of proliferation) or after 21 days (assessment of survival).

### Tissue processing

Mice were anesthetized, trans-cardially perfused with saline and the brains have been dissected. The right hemispheres were further dissected and hippocampal specimens were stored in − 80°C for RNA isolation. The left hemispheres were post-fixed by 4% paraformaldehyde (158127, Sigma) overnight at 4oC. They were stored in cryoprotective medium (15% sucrose/7,5% gelatine) at − 80°C, until they processed for coronal sections. Coronal sections of 40 μM were cut in the dorsoventral axis of hippocampus (from bregma -4 mm to -1mm).

### Immunohistochemistry

Cryosections were permeabilized by immersion in ice-cold acetone at 20°C for 5 min and washed with 0.1% Triton X-100 in 1xPBS for 15 min and 0.3% Triton X-100 in 1xPBS for 30 min, then blocked for 1hr in 10% donkey serum (S30, Millipore, Burlington, MA, USA) containing 0.1% Triton X-100 in 1xPBS and 0.1% BSA, and incubated overnight at 4°C with the above primary antibodies (Table 1). Slides were then washed and incubated with the appropriate fluorochrome-labeled secondary antibodies at room temperature (Table 1). Cell nuclei were visualized with Hoechst (1:10.000, H3570, Invitrogen, Carlsbad, CA, USA). Slides were covered with VECTASHIELD® Antifade mounting medium (VECTOR, Newark, CA, USA) and images were photographed via confocal microscopy (SP8 Leica, Wetzlar, Germany).

**Table 1:** Summary of the *in vivo* results.

| Age | Markers | WT vs p75NTR ko | WT vs 5xFAD | 5xFAD vs 5xFAD/p75NTR ko |
| --- | --- | --- | --- | --- |
| 2 months old | BrdU/Sox<sub>2</sub> | ↓ | ↑ | ↓ |
|  | Dcx | ↑ | ↑ | ↓ |
| 4 months old | BrdU/Sox<sub>2</sub> | ↓ | ↓ | ↓ |
|  | Dcx | Not changed | ↓ | Not changed |
| Age | Markers | WT vs p75NTR ko | WT vs 5xFAD | 2mo WT vs 6mo WT |
| 6 months old | BrdU/Sox<sub>2</sub> | Not changed | Not changed | ↓ |
|  | Dcx | Not changed | Not changed | ↓ |

For double labelling and the detection of BrdU-labeled nuclei, specimens have been previously incubated in 2N HCl at 37°C, followed by two rinses with PBS before blocking step.

### Cell counts and quantification

Cell counts and quantification are based on a modified unbiased stereology protocol. Seven out of every 10 adjacent sections were chosen (covering the whole DG area of the hippocampus) and processed for immunohistochemistry. The number of BrdU+, Sox2+, Dcx+, NeuN+ and caspase+ cells was then counted under ×40 magnification under a fluorescent microscope (Leica sp8) at the area of granular cell layer and SGZ of a total of 7 sections and the average number of cells was estimated. The mean was then multiplied with the total number of sections (75 per mouse) to estimate the total number of cells per DG.

### Generation and Culture of Human Neural Stem Cells (hNSCs)

iPSCs were reprogrammed from skin fibroblasts of two healthy human lines (SFC856-03-04, SFC841-03-01) that were kindly provided by Dr Μ. Z. Cader and used for the generation of neural progenitor cells, as previously described in (Reinhardt et al., 2013). Briefly, iPSCs colonies were cultured on mouse embryonic fibroblasts (MEFs), in a medium that consisted of DMEM-F12 (21331-020, Gibco, Carlsbad, CA, USA), 20% (v/v) Knockout Serum Replacement (10828028, Gibco), 1% non-essential amino acids (NEAA; 11-140-050, Gibco), 1% penicillin/streptomycin (PAA; 15140122, Gibco), L-Glutamine (A2916801, Gibco), 2-Mercaptoethanol (31350010, Gibco) supplemented with 5 ng/ml FGF2 (100-18C, Peprotech, Cranbury, NJ, USA). Next, iPSC colonies were detached from MEFs with 2 mg/mL collagenase IV (C1764, Sigma) and resuspended in the same medium without FGF2, supplemented with 1 μM Dorsomorphin (ab120843, Abcam, Cambridge, UK), 3 μM CHIR99021 (SML1046, Sigma), 10 μM SB-431542 (72232, Stem Cell Technologies, Vancouver, BC, Canada) and 0.5 μM Purmorphamine (72202, Stem Cell Technologies). Embryoid bodies (EBs) were formed and medium was changed to N2B27 medium (1:1 Neurobasal (21103-049, Gibco) and DMEM-F12 medium, supplemented with N2 supplement (17502048, Gibco) and B27 supplement lacking vitamin A (12587010, Gibco), and 1% penicillin/streptomycin supplemented with the aforementioned small-molecules. On day 4, dorsomorphin and SB-431542 were removed, whereas 150μM L-Ascorbic acid (A4544, Sigma) was added to the medium. On day 6, the spheres were cut into smaller pieces and plated on Matrigel-coated plates (354263, Corning, NY, USA). When confluent, cells were split via treatment with accutase (A6964, Sigma). The identity of the cells was verified via immunocytochemistry for NESTIN (151 NB100-1604, BioTechne, Minneapolis, MN, USA). The generated NSCs can be expanded and cryopreserved enabling long-term, repetitive studies.

### Immunoprecipitation and Immunoblotting

Cells were suspended in Pierce™ IP Lysis Buffer (87788, Thermo Fischer Scientific, Waltham, MA, USA) supplemented with protease inhibitors (539138, Calbiochem, Burlington, MA, USA) and phosphatase inhibitors (524629, Calbiochem). Lysates were pre-cleared for 1h with protein G-plus Agarose beads (sc-2002, Santa Cruz Biotechnology, Dallas, TX, USA) and immunoprecipitated with p75NTR antibody overnight at 4 °C. Protein G-plus agarose beads were incubated with the lysates for 4h at 4 °C via gentle shaking. Beads were collected via centrifugation, re-suspended in 2× SDS loading buffer and subjected to Western blot analysis against TRAF6 antibody. For immunoblot (IB) analysis, the beads were suspended in sodium dodecyl sulfate-loading buffer and separated through SDS-PAGE. Proteins were transferred to nitrocellulose membranes and blotted with the corresponding antibodies for RIP2, p75NTR, TRAF6 and Actin (Table 1). Immunoblots were developed using the ECL Western Blotting Kit (ThermoFisher Scientific), and Image analysis and quantification of band intensities were performed with ImageJ Software. For the immunoprecipitation assay, the analysis was derived from the immunoprecipitated fraction relative to the total fraction.

### Preparation of Aβ Oligomers

Amyloid-β (1–42) peptide was purchased from AnaSpec (AS-20276, AnaSpec, Fremont, CA, USA) and prepared according to manufacturer’s instructions. For Aβ treatment, Aβ oligomers were prepared according to previously described protocols (Kokkali et al., 2024), (S. Li et al., 2011) and they were diluted in DMEM at the specified concentrations. Human NSCs were treated with 10μΜ for 48 hours.

### Cell Tox Assay

After 24h of treatments, we used the CellTox™ Green Cytotoxicity Assay kit (G8742, Promega Corporation, Madison, WI, USA) to assess the survival of hNSCs in the presence or absence of p75NTR inhibitor MC-192 (2.5 ng/mL, ab6172, Abcam). AD pathology was mimicked by treating the human NSCs with Aβ-amyloid peptides (10μΜ of Aβ1-42 oligomers) and after 24hours we used cell tox assay for another 24h. Dead cells were then counted with a fluorescent microscope at 485–500nm Ex. CellTox assay reagents and Hoescht (1:10,000, H3570, Invitrogen) were added to each well for 30 min, and cells were imaged using a fluorescent microscope (Zeiss AXIO Vert A1, Zeiss, Oberkochen, Germany). Positive cells for cell tox reagent were normalized to reflect the total number of cells per image.

### 5-bromo-2′-deoxyuridine (BrdU) assay in hiPSCs-derived NSCs

iPSCs-derived NSCs were cultured on Matrigel for 24h with or without treatment of p75NTR inhibitor (ab6172, abcam MC-192, 2,5ng/ml). After 24h the cells were pulsed with 1μM BrdU for 4 hours and fixed with 4% PFA for subsequent immunostaining for BrdU and Hoechst for nuclear labeling.

### Isolation of RNA & Sequencing

Total RNA from biological triplicates (hippocampal specimens of p75NTR ko, 5xFAD, 5xFAD/p75NTR ko and WT mice of 2 months old, that have been dissected after the BrdU injections) was extracted using Trizol reagent (15596018, Thermo Scientific) as per the manufacturer’s protocol. The quantity and quality of RNA samples were analyzed using Agilent RNA 6000 Nano kit with the bioanalyzer from Agilent. RNA samples with RNA integrity number (RIN)>7, were used for library construction using the 3′ mRNA-Seq Library Prep Kit FWD for Illumina (QuantSeq-LEXOGEN, Vienna, Austria) as per the manufacturer’s instructions. Amplification was controlled for obtaining optimal unbiased libraries across samples by assessing the number of cycles (14) required by qPCR. Indexes used are shown in Table 2. DNA High Sensitivity Kit for bioanalyzer was used to assess the quantity and quality of libraries, according to the manufacturer’s instructions (Agilent, Santa Clara, CA, USA). Libraries were multiplexed and sequenced on an Illumina Nextseq 2000 at the genomics facility of IMBB FORTH according to the manufacturer’s instructions.

**Table 2:**
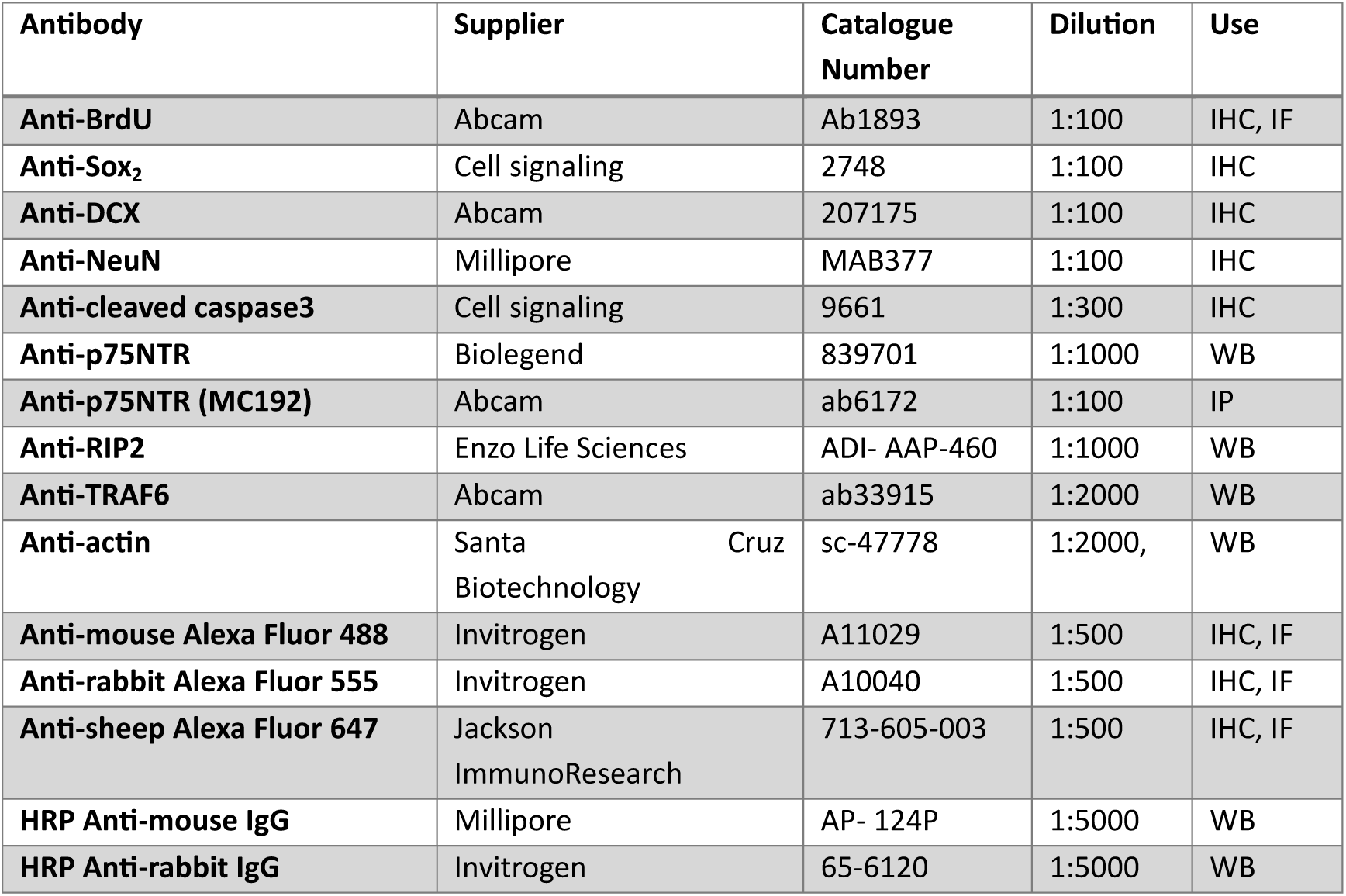
List of primary and secondary antibodies used in Immunohistochemistry-Immunofluorescence and Western blot assay.

**Table 3:** Indexes used for RNA-seq library preparation and differential expression analysis.

|  |  |
| --- | --- |
| WT 1 | Lex_i712_0089 |
| WT 2 | Lex_i712_0090 |
| WT 3 | Lex_i712_0091 |
| 5xFAD 1 | Lex_i712_0092 |
| 5xFAD 2 | Lex_i712_0093 |
| 5xFAD 3 | Lex_i712_0094 |
| p75NTR ko 1 | Lex_i712_0095 |
| p75NTR ko 2 | Lex_i712_0096 |
| p75NTR ko 3 | Lex_i712_0071 |
| 5xFADp75NTR ko 1 | Lex_i712_0072 |
| 5xFADp75NTR ko 2 | Lex_i712_0048 |
| 5xFADp75NTR ko 3 | Lex_i712_0036 |

### Differential Expression and GO enrichment analysis

The quality of the raw sequences in the output FASTQ files was assessed with the FastQC software [www.bioinformatics.babraham.ac.uk/projects/fastqc/]. Reads were aligned to the mouse (mm10) genome using the Hisat2 aligner (parameters used: hisat2 -p32 -x $REFERENCE_GENOME -q fastq/ $FILE_ID.fastq -S $FILE_ID.sam --score-min L,0,-0.5) (D. Kim et al., 2019). The BAM files were sorted by genomic coordinates and indexed using samtools (H. Li et al., 2009). Htseq-count was utilized to summarize reads at the gene level (parameters used: htseq-count -f bam -s yes -i gene_id bam $FILE_ID.bam $REFERENCE_ANNOTATION > $COUNTS_DIR/NGS$FILE_ID) (Anders et al., 2015). Differential expression analysis (DEA) was conducted using EdgeR (Chen et al., 2025). Two designs were employed, both accounting for gender differences between the mice (gender included as a covariate). The first design also included disease and genotype as covariates (∼ disease + genotype + gender), with healthy WT mice treated as the reference. The second design accounted for mouse line as a group covariate (∼ group + gender, groups: WT, 5xFAD, p75NTR ko, 5xFAD/p75NTR ko), with WT mice as the reference. Differentially expressed genes (DEGs) were identified using a significance threshold of adjusted p-value (padj) < 0.05. Enrichment analysis was performed using the Metascape web tool (enrichment p-value set to 0.05) (Zhou et al., 2019). Gene Set Enrichment Analysis (GSEA) tool was also used, to analyze the functional changes in gene expression across genotype (p75NTR ko vs WT) and between 5xFAD and 5xFAD/p75NTR ko samples, as specified by Broad GSEA package using default parameters (Subramanian et al., 2005). GO-BP pathways were tested (https://www.gsea-msigdb.org/gsea/msigdb/mouse/genesets.jsp?collection=GO:BP) against a ranked lists of descending LFCs for both contrasts, and we selected some positive hits for display. The volcano and bubble plots were created in R with custom in-house scripts (available upon request).

### Reverse Transcriptase PCR & Quantitative PCR

cDNA was synthesized by the total RNA, using the High-Capacity cDNA Reverse Transcription kit (4368814, Thermo Fisher) according to the supplier protocols. Primers for Ngfr, were designed using the NCBI Primer BLAST software (Forward:AGAGAAACTGCACAGCGACA, Reverse:CCATCACCCTTGAGGGCTTG), to detect an area inside the coding sequence (CDS) of the gene. More specifically, the primer pair was designed as such to amplify specifically the area between exons V and VI (between TMD – Transmembrane Domain and DD – Death Domain). Primers for mouse GAPDH, were also designed to be used as a control sample (Forward: ATTGTCAGCAATGCATCCTG, Reverse: ATGGACTGTGGTCATGAGCC). To run the quantitative RT-PCR, we used 1 μL of cDNA (10 ng/μL) and the KAPA SYBR Fast kit (KK4601, Sigma) according to the supplier’s instructions. The cycling program consisted of 20 s at 95 °C, followed by 40 cycles of 95 °C for 3 s and 60 °C for 30 s on a StepOne Real-Time PCR System (Thermo Fisher Scientific). After the completion of qPCR, a melt curve of the amplified products was performed. The housekeeping gene GAPDH was used to normalize the expression levels between the different conditions. Data were collected and analyzed using the StepOne Software v2.3 (Thermo Fischer Scientific).

### Statistical Analysis

All values are expressed as the mean ± SEM. Student’s t-test was used for the comparison of two groups, and one-way or two-way ANOVA were used for multiple group comparisons. A p<0.05 was considered to mark statistical significance. Statistical analysis was performed using GraphPad Prism 7 (GraphPad Software Inc., San Diego, CA, USA).

## Data availability

The RNA-seq datasets generated and analyzed during the current study are available in the GEO repository GSE296390. All materials are available upon request. Expanded view data, supplementary information, appendices are available for this paper.

## Acknowledgements

We thank Dr Sebastian Thieme from TU of Dresden for providing us the p75 fl/fl mice (Ngfrtm1a^(EUCOMM)Wtsi^), Alexandros Tsimpolis (PhD student at the University of Crete) for performing qRT-PCR and Dr Μ. Z. Cader for providing the two hiPSC lines.

## Author Contributions

M.A.P.: conception and design of the study, acquisition and interpretation of the data, drafting the text, MΑP, KC, MP: *in vivo* experiments, *in vitro* experiments, analysis of results, I.C.: conception and design of the study, drafting the manuscript, supervision, funding, E.T., K.M., M.L.: RNA-seq analysis

## Institutional review board statement

All procedures were performed under the approval of Veterinary Directorate of Prefecture of Heraklion (Crete) and carried out in compliance with Greek Government guidelines and the guidelines of FORTH ethics committee and were performed in accordance with approved protocols from the Federation of European Laboratory Animal Science Associations (FELASA) and Use of Laboratory animals [License number: EL91-BIOexp-02), Approval Code: 360667, Approval Date: 29/11/2021 (active for 3 years)]. All research activities strictly adhered to the EU adopted Directive 2010/63/EU on the protection of animals used for scientific purposes.

---

**Figure EV1.**
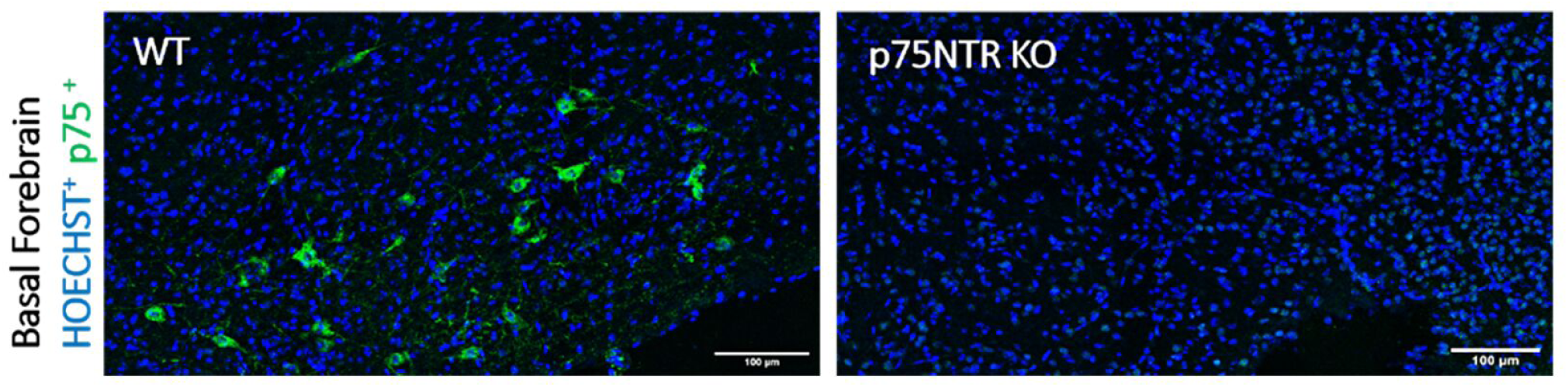
p75NTR expression in 2 months old p75NTR ko mice in the Basal Forebrain. p75NTR expression at the basal forebrain of wild type and knock out mice. Coronal sections 40μm from 2-months-old mice after immunostaining with anti-p75 antibody and Hoechst for nuclear staining (WT, wild type; KO, knock out, scale bar: 100μm).

**Figure EV2.**
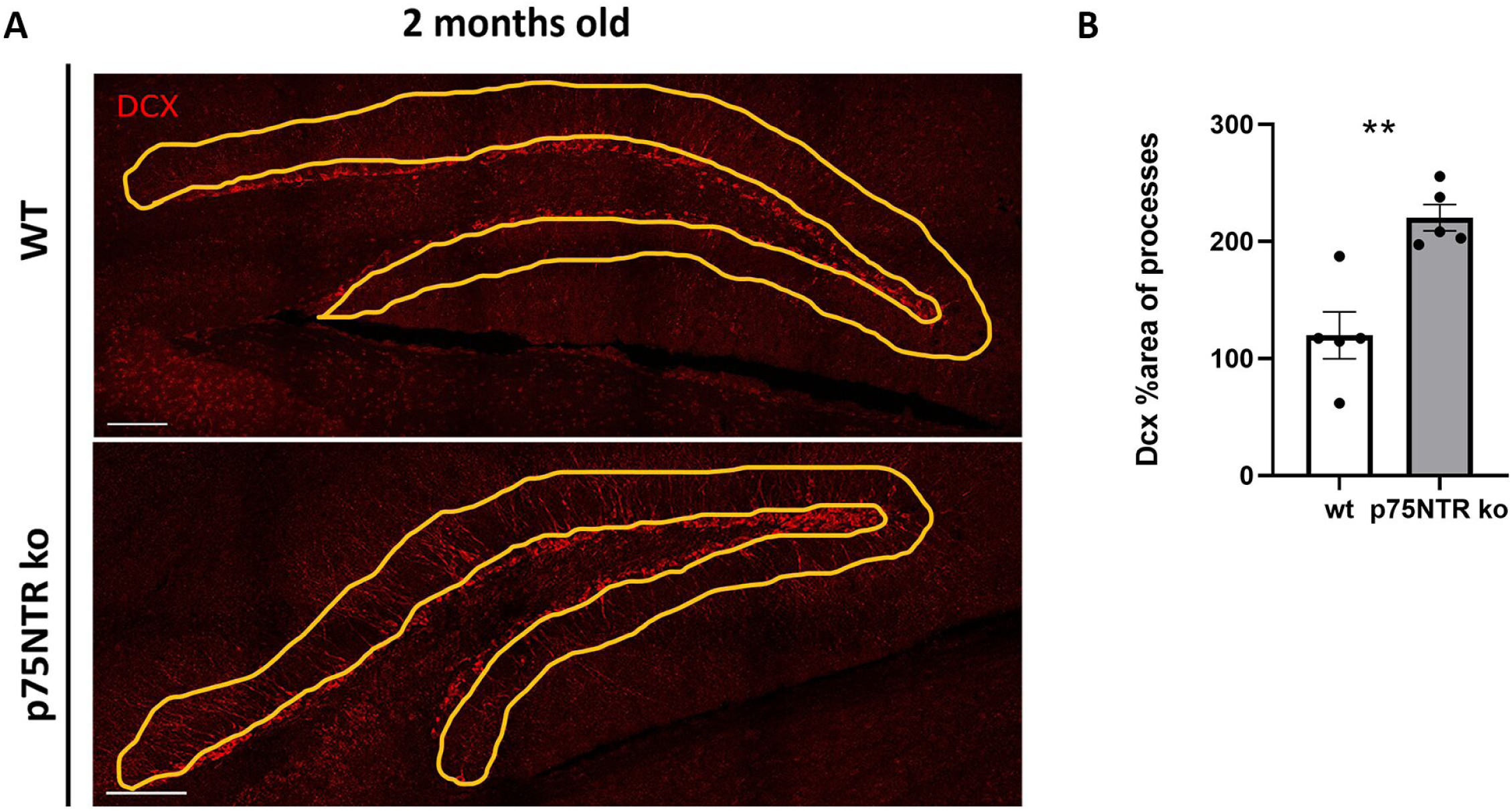
Dcx area of processes in 2 months old WT vs p75NTR ko mice. (Α) Coronal section of DG from 2-months-old wild type & p75NTR ko mice. Image depicts Dcx (red) immunostained immature neurons. (Β) Quantification of Dcx area of processes in p75NTR ko & wild type mice of 2 months old (n=5 ko, n=5 wt, unpaired t-test, **p<0,005). DcX, doublecortin.

**Figure EV3.**
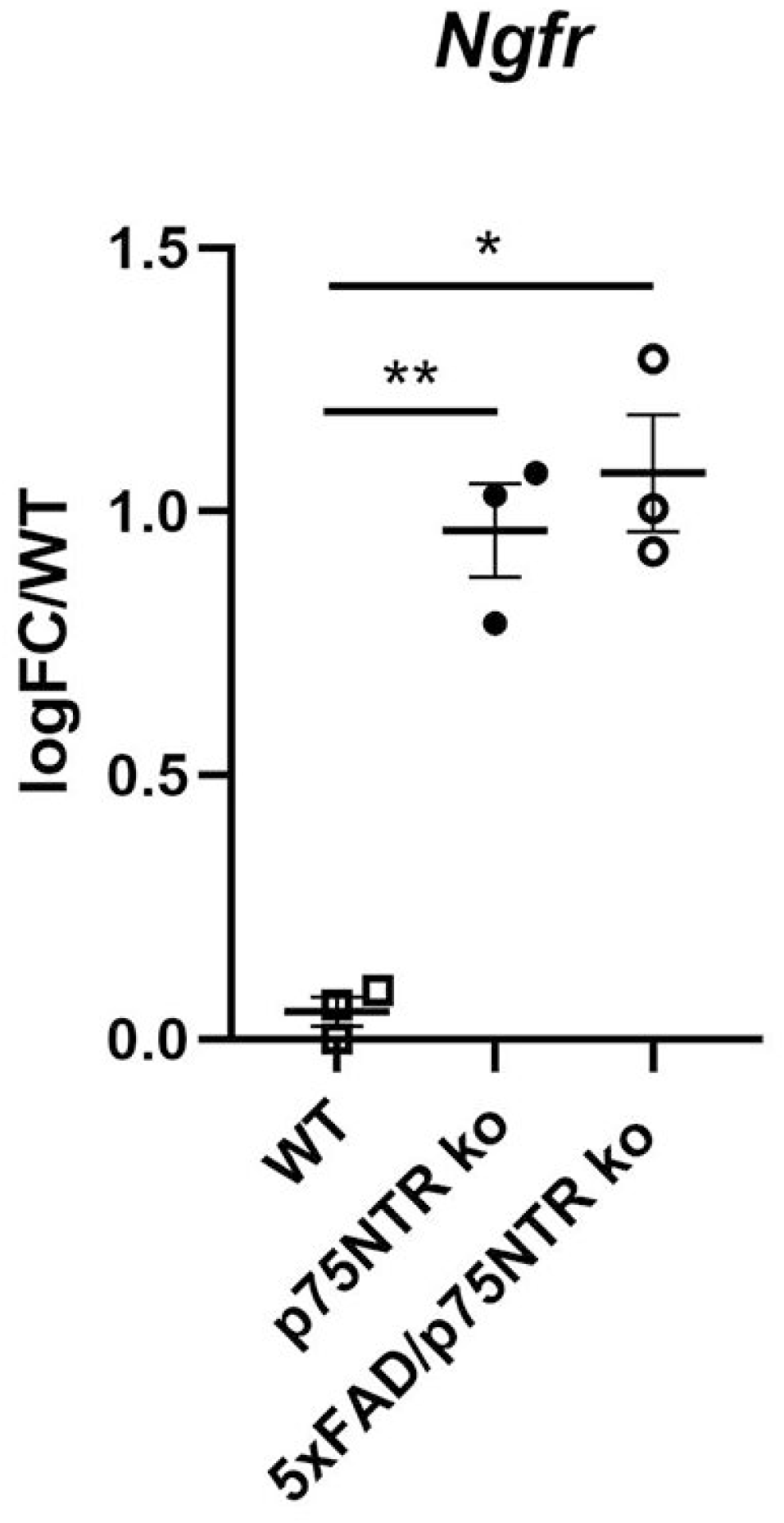
qRT-PCR was performed in RNA samples. Total RNA from biological triplicates (hippocampal specimens of p75NTR ko, 5xFAD/p75NTR ko and WT mice of 2 months old, that have been dissected after the BrdU injections) was extracted using Trizol reagent. qRT-PCR was performed in RNA samples and the results are expressed as the logarithmic fold change (logFC) normalized to the WT control. p75NTR exon III deletion, strongly upregulates Ngfr mRNA expression levels on p75NTR ko, 5xFAD/p75NTR ko. (n = 3 for each mouse model. * p < 0.05, ** p < 0.01; unpaired t-test).

**Figure EV4.**
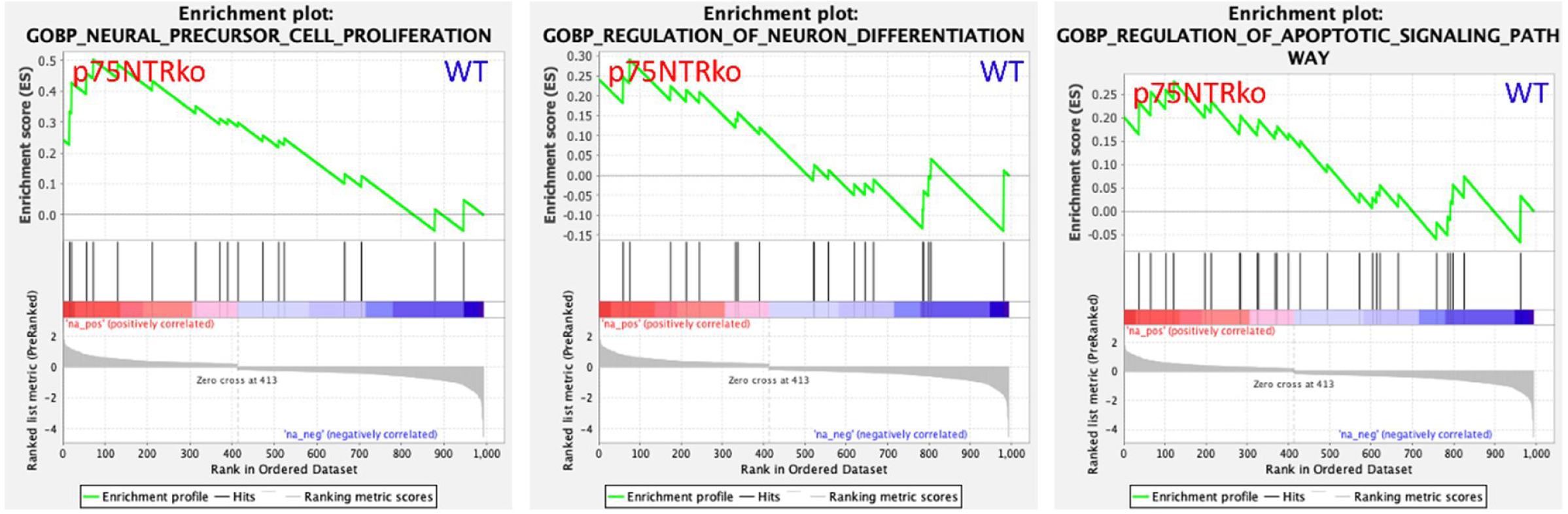
GSEA of RNA-seq data showing enrichment of selected GO_BP categories genes in p75NTR ko vs WT mice. GO for the Neural Precursor Cell Proliferation, GO for the regulation of neuron differentiation and GO for the regulation of apoptotic signaling pathway. Gene sets were obtained from MSigDB. Normalized enrichment scores (NES) (y-axis) and false discovery rate (FDR) were calculated by GSEA based on 1,000 permutations.

**Figure EV5.**
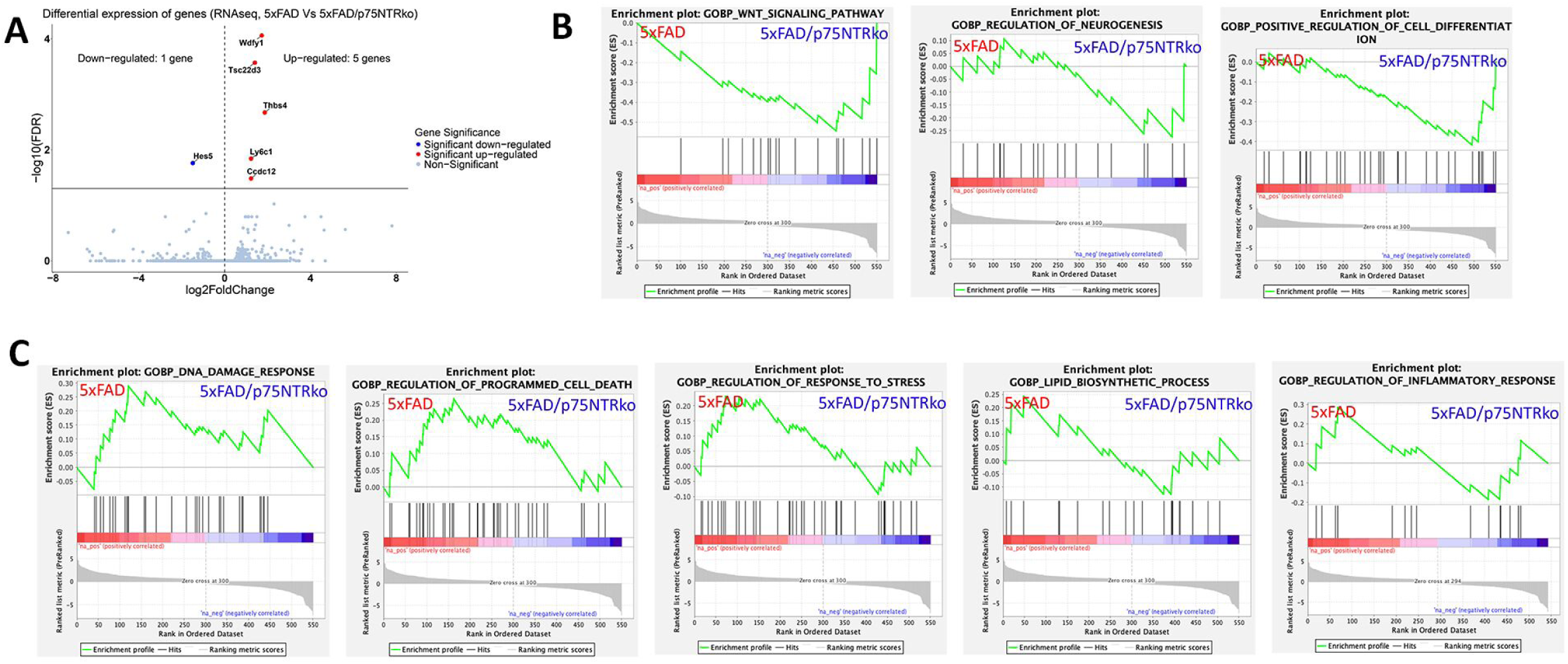
Volcano plot and GSEA analysis of RNA-seq data. Volcano plot and GSEA analysis of RNA-seq data derived from the comparison between 5xFAD vs 5xFAD/p75NTR ko mice. (A) Volcano plot showing the number of Up (5) & Down (1) regulated genes (p adj < 0.05). (B) GSEA analysis of RNA-seq data showing enrichment of selected GO_BP categories genes in 5xFAD vs 5xFAD/p75NTR ko mice (functions which are enriched in 5xFAD/p75NTR ko samples). (C) GSEA analysis of RNA-seq data showing enrichment of selected GO_BP categories genes in 5xFAD vs 5xFAD/p75NTR ko mice (functions which are enriched in 5xFAD samples). Gene sets were obtained from MSigDB. Normalized enrichment scores (NES) (y-axis) and false discovery rate (FDR) were calculated by GSEA based on 1,000 permutations.

